# SVhound: Detection of future Structural Variation hotspots

**DOI:** 10.1101/2021.04.09.439237

**Authors:** Luis F Paulin, Muthuswamy Raveendran, R. Alan Harris, Jeffrey Rogers, Arndt von Haeseler, Fritz J Sedlazeck

## Abstract

Recent population studies are ever growing in size of samples to investigate the diversity of a given population or species. These studies reveal ever new polymorphism that lead to important insights into the mechanisms of evolution, but are also important for the interpretation of these variations. Nevertheless, while the full catalog of variations across entire species remains unknown, we can predict which regions harbor additional variations that remain hidden and investigate their properties, thereby enhancing the analysis for potentially missed variants.

To achieve this we implemented SVhound (https://github.com/lfpaulin/SVhound), which based on a population level SVs dataset can predict regions that harbor novel SV alleles. We tested SVhound using subsets of the 1000 genomes project data and showed that its correlation (average correlation of 2,800 tests r=0.7136) is high to the full data set. Next, we utilized SVhound to investigate potentially missed or understudied regions across 1KGP and CCDG that included multiple genes. Lastly we show the applicability for SVhound also on a small and novel SV call set for rhesus macaque (*Macaca mulatta*) and discuss the impact and choice of parameters for SVhound. Overall SVhound is a unique method to identify potential regions that harbor hidden diversity in model and non model organisms and can also be potentially used to ensure high quality of SV call sets.

## Introduction

The advent of next generation sequencing has enabled us to characterize genomic variations between and within species on an unprecedented scale (Lappalainen et al. 2019; Goodwin et al. 2016). This has produced various novel insights based on sequence complexity and previously underestimated genomic variability between individuals within the same species (Sudmant et al. 2015). Since then, reports have described an ever-increasing number of novel genomic variations and their associated allele frequency estimates (Sedlazeck et al. 2020; Collins et al. 2020; Sudmant et al. 2015; Ebert et al. 2021; Audano et al. 2019; Warren et al. 2020). These findings are important for many fields in research and clinical applications, ultimately providing a better understanding of phenotype to genotype relationships (Lappalainen et al. 2019; Mahmoud et al. 2019; Ho et al. 2020).

Over the past years, genomic studies emerged targeting even higher sample numbers to obtain deeper insights into allele frequencies and diversity (genomic variation) among humans or other species (Collins et al. 2020; Sedlazeck et al. 2020; Sudmant et al. 2015; Abel et al. 2018). One of the spearheading projects in the past years was the 1,000 Genomes Project (1KGP), which cataloged single nucleotide variations (SNV) and structural variations (SV) among 2,504 individuals from different ethnicities around the world (Sudmant et al. 2015). While it is clear that the 1KGP catalog is incomplete, it is still one of the most valuable datasets and it is widely used as control data (Sudmant et al. 2015). More recent initiatives such as gnomadSV investigated the presence of SVs across 14,891 human genomes and thus deepened our knowledge of human genome diversity (discovering ~445k SVs) and allele frequencies that are important for multiple aspects (Collins et al. 2020), such as ranking and annotating variations or identifying population structure. However, even larger studies are underway (e.g. Topmed (Taliun et al. 2019), CCDG (Abel et al. 2018)) that will identify many new SNVs/SVs in presumably healthy individuals and lead to even more robust ethnicity specific allele frequencies and also to a better understanding of variability with respect to diseases.

The detection of genomic variations is often promoted by technological and methodological advances in computational methods (Mahmoud et al. 2019; Wenger et al. 2019). As an example, microarrays enabled the first identification of so-called large copy number variations (CNV), in the range of kbp to Mbp, at scale (Sebat 2004). Subsequently, short read sequencing technologies (whole exome or whole genome sequencing) detected these large alterations and SNVs simultaneously. Many developments in computational methods led to a better characterization of large events (e.g. CNV of multiple kbp) and identification of even more complex structural variations (Mahmoud et al. 2019). The continuous advance of better benchmark datasets (e.g. GIAB (Zook et al. 2020)) and software will lead to many newly identified variations in currently hard to assess regions (e.g. dark regions) of the genome.

Despite these developments and the increased number of studies sequencing hundreds to thousands of humans, we still expect an unknown number of undetected genomic variations including rare or even common alleles. This is especially true for ethnicities that have not yet been extensively sequenced (e. g. non-European) (Audano et al. 2019).

Thus, the questions arise: which genomic regions carry novel yet undetected variations in our enlarged datasets? Can we predict such genomic regions based on existing sequencing data, and if so where are these regions located in the genome and what else can we learn about the mechanisms generating SVs?

To address these questions, we utilized large genomic SV datasets from the 1KGP (Sudmant et al. 2015) and CCDG (Sedlazeck et al. 2020) cohorts and applied a population genetic approach that computes the likelihood to observe novel genomic variations, if we had sequenced more individuals. To this end we developed SVhound, which scans the genome for regions of hidden diversity. In the following we demonstrate the predictive power of SVhound based on the analysis of the 1KGP dataset. Next, we applied SVhound to the CCDG cohort composed of a collection of 19,652 human samples (Sedlazeck et al. 2020). Finally, SVhound is applied to uncover regions of undetected genomic variability in genomes from 150 rhesus macaques (Macaca mulatta), an important model species for human diseases and evolutionary studies. Currently, little is known about SVs in rhesus macaques (Brasó-Vives et al. 2020; Thomas et al. 2020). SVhound introduces a novel prediction framework to identify genomic regions that are lacking genotypes from current large-scale sequencing and studies the properties of these regions and their potential role. Finally, we provide an easy to use R package freely available at https://github.com/lfpaulin/SVhound.

## Results

### Statistical identification of highly variable genomic regions in the human population

Here we present SVhound, a tool to predict regions where additional Structural Variation (SV, defined as genomic variation greater 50bp) can be expected if more genomes were sequenced. In short, SVhound partitions a genome into non-overlapping windows with a user-defined length. For each window, SVhound counts the number of different SV-alleles that occur in a sample of *n* genomes. Based on the number of different SV-alleles, SVhound then estimates the probability of observing a new SV-allele (see Methods).

**Figure 1A** exemplifies this for three windows and a sample of *n*=100 genomes. In windows *w_1_, w_2_, w_3_*, we detected *k*=3, 5, 2 SV-alleles, leading to diversity parameter estimatesθ(*w*_1_) = 0. 430, θ(*w*_2_) = 0. 948, θ(*w*_3_) = 0. 204 and the probabilities to find a new SV-allele in the respective windows, if an additional genome or sequence from the respective window is sequenced, equal *p*_*new*_ (*w*_1_) = 0. 00430, *p*_*new*_ (*w*_2_) = 0. 009390, *p*_*new*_ (*w*_3_) = 0. 00205.

**Figure 1:**
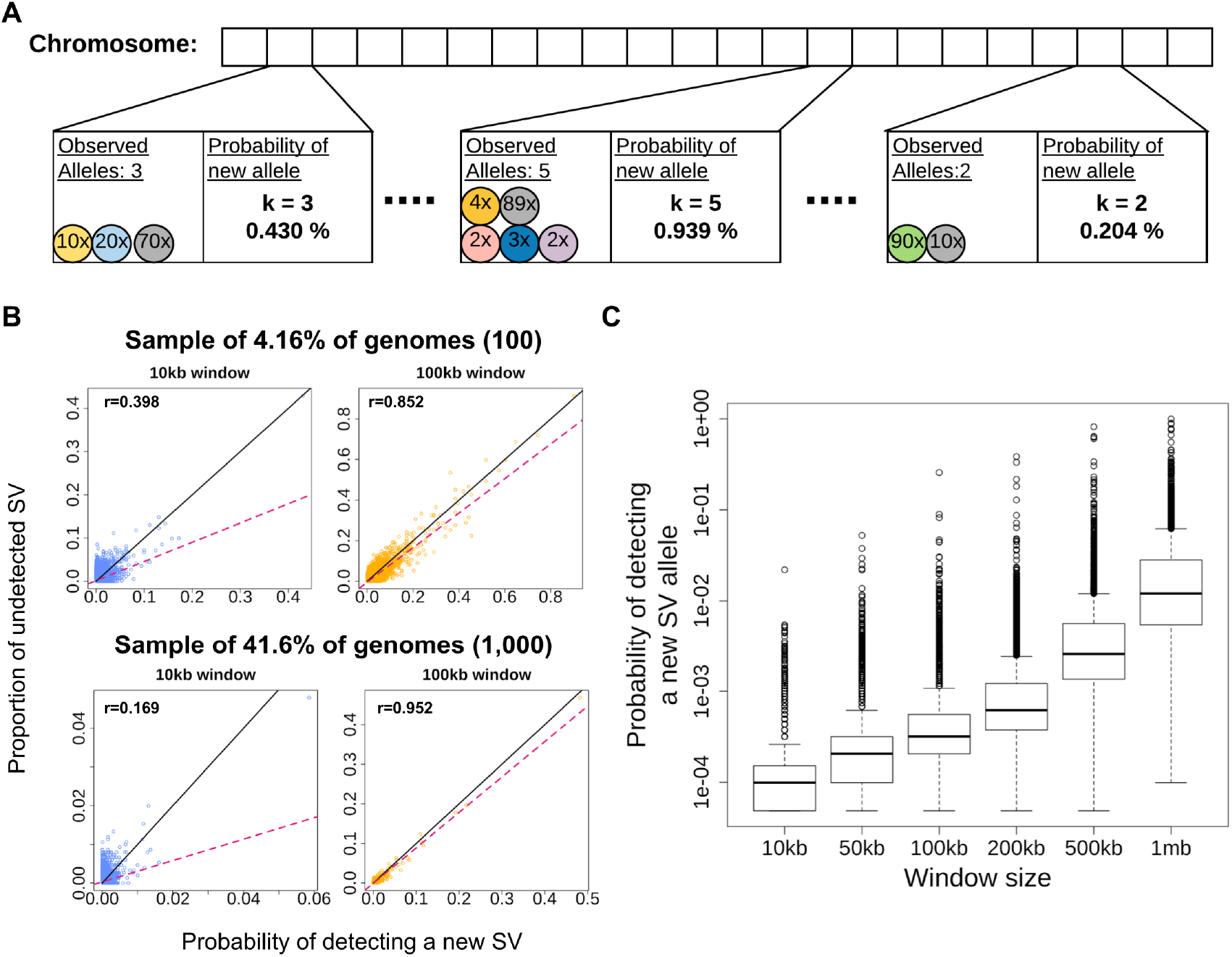
Overview and evaluation of SVhound based on 1000 genomes data set. **A)** Computing the probabilities of detecting new SV-alleles in a window. First, the chromosome is divided into non overlapping windows. For each window the number of distinct observed SV-alleles is counted and the diversity parameter is estimated equation 2 (see methods). Finally, the probability of detecting a new SV-allele (*p*_*new*_) for each particular window is computed using equation 3 (see methods). **B)** Scatterplots showing predictive power (correlation) between *p*_*new*_ and the fraction of undetected SV for a 10kb and 100kb window and two sample sizes 100 genomes (top panels) and 1,000 genomes (bottom panels), sub-sampled from the 1KGP data. The x-axis shows the prediction made by SVhound (probability of new SV-allele, *p*_*new*_) and the y-axis shows the proportion of undetected SV-alleles in the non-sampled individuals (*f*_*undetected*_). Note that regardless of sample size, SVhound performs better in the 100kb window when comparing both window sizes. **C)** Distribution of the probabilities of detecting a new SV-allele (*p*_*new*_) for different window sizes.

To further investigate the power of SVhound to predict new SV-alleles and to study the influence of the window-length, we randomly selected 50 (2.00%), 100 (4.00%), 500 (19.97%) and 1000 (39.34%) human genomes from the 2,504 genomes of the 1KGP (1000 Genomes Project Consortium et al. 2015) and varied the window size (5, 10, 50, 100, 200, 500 and 1000 kb). For each of the 28 combinations of window size and sample size we compared the *p*_*new*_ estimates with the fraction *f*_*undetected*_ of SV-alleles that do not occur in the random sample but were observed in the full 1KGP data (see Methods).

**Figures 1B** display the association between *p*_*new*_ and *f*_*undetected*_ for a sub-sample size of n=100 (**Figure 1B top panel**) and a sub-sample size of n=1000 genomes (**Figure 1B bottom panel**) and window lengths of 10kb and 100kb, respectively. We observed that the window size had a bigger impact on the performance of *p*_*new*_; for example the correlation coefficient (r) for 10kb window is r=0.3976 and r=0.1698 for 100 and 1,000 genomes respectively (**Figure 1B top panel**), while for 100kb window the performance of SVhound greatly improves with r=0.8519 for 100 genomes and r=0.9524 for 1,000. We also noticed that the sample size only improved the correlation coefficient for window sizes of at least 50kb. The scatterplots of the 28 window-sample size combinations are shown in **Supplementary Figure 1**.

While the above analysis was based on one simulation, we performed 100 simulations for each of the 28 parameter combinations. **Supplementary Figure 3** and **4** show the distribution of the correlation coefficients, the coefficients of determination (*r*^2^) and the slopes for the 100 simulations and the observations exemplified in **Figure 1B** are corroborated.

**Supplementary Table 1.1** shows the average correlation coefficients for the 100 simulations for each of the 28 window-sample size combinations. If the window size is large and the sample size is large then we observe a high correlation between *p*_*new*_ and *f*_*undetected*_. Since large windows harbor more SV-alleles, the infinite allele assumption is almost met and thus the predictions improve. For short windows the model assumptions (infinitely many alleles) are more likely violated and thus the correlation is weaker. But not only the correlation is high for large windows, also the slope of the regression line approaches one with increasing sample size and window length (Table 1.2). This indicates that *p*_*new*_ is indeed a good predictor of *f*_*undetected*_.

We note, that with increasing window length *p*_*new*_ increases (see also **Figure 1C**), while the increase in sample size has the opposite effect (**Supplementary Figure 5)**. This can also be explained with the infinite allele assumption almost being met and thus the probability to find new SV-alleles increases. With increasing window length the chances also increase to find many SV-alleles that occur exactly once, high numbers of such singletons will increase the diversity parameter, θ, and subsequently *p*_*new*_ (see methods). However, with larger window sizes the resolution and thus the genomic location of the predicted additional SV-alleles is reduced.

### Identification of polymorphic candidate regions across 2,504 human genomes from the 1,000 genome project

We applied SVhound to the 2,504 genomes of 1KGP SV calls to identify likely regions (loci) with undetected SV variations. Based on the previous analysis we opted for a window size of 100kb. The human genome was then partitioned into 18,397 windows and we analyzed the top candidate loci, representing 1% of the windows with the highest probability of detecting a new SV-alleles (*p*_*new*_ ≥ 0. 34%). **Figure 2A** shows the probability distribution of detecting a new SV-allele for each window. The red dots mark the top 1% (188) windows with the highest *p*_*new*_ (here thereafter candidate windows). The remaining windows with *p*_*new*_ < 0. 34% (considered as background noise) are gray. We observed an outlier on chromosome 15 with a *p*_*new*_ = 25. 77%.

**Figure 2.**
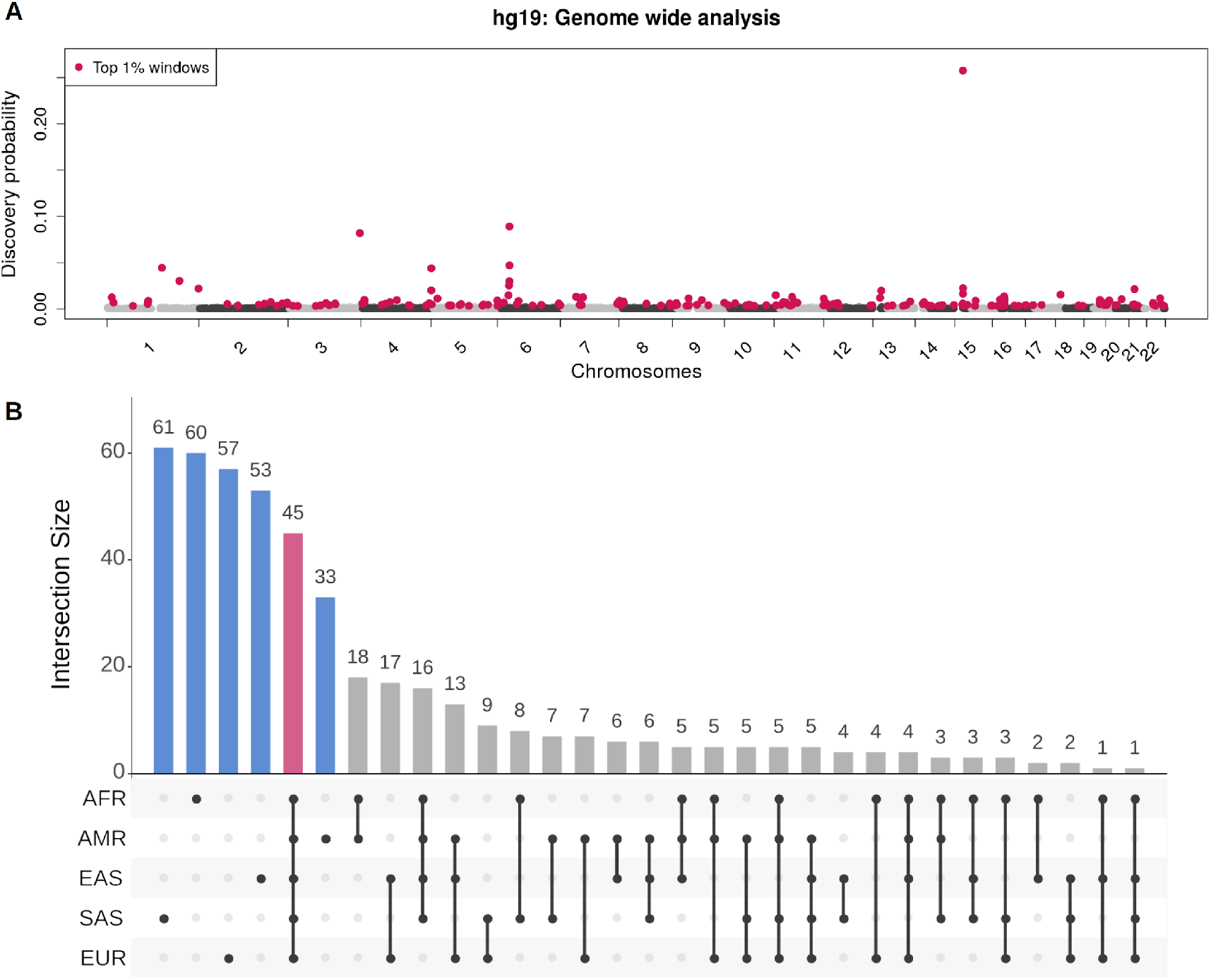
**A)** Genome wide distribution of *p*_*new*_ for the 2504 genomes (100 kb window) from the 1KGP data set. Red dots show the 188 candidate windows (*p*_*new*_ ≥ 0. 34%) along the 22 human autosomes (hg19), gray/black (alternating shades by chromosome) dots display the *p*_*new*_ for the remaining windows. Please, note the window on chromosome 15 with a *p*_*new*_ = 25. 77%, contains two olfactory receptor proteins, four olfactory receptor pseudogenes, multiple CNVs and an LINE1 insertion. **B)** Distribution of 468 candidate windows when decomposing the 1KGP data set into the five super-population: African, AFR; Admixed American, AMR; European, EUR; East Asian EAS; South Asian, SAS. The black dots below each bar display the occurrences of the candidate windows in the ethnic groups. Ethnicity specific windows, i.e present in one ethnic group are blue, ubiquitous windows are red.

We were particularly interested where in the human genome the 188 candidate windows occur. First, we investigated whether these candidate windows are identified only in intergenic regions or if these windows are actually preferentially selected for certain genes (e.g. immune response). We found 107 candidate windows that overlapped with protein coding genes (204, **Supplementary table 2**), 148 overlapping non coding genetic elements (**Supplementary figure 6**) and 24 windows in intergenic regions. To understand the biological role of the 204 genes we performed an enrichment analysis with PANTHER (Mi et al. 2019), and found enrichment for biological processes related to: cellular detoxification of nitrogen compound, xenobiotic catabolic process, interferon-gamma-mediated signaling pathway, regulation of immune response and sensory perception of smell (**Supplementary table 2** and **3**). The outlier we observed on chromosome 15 contains two olfactory receptor proteins and four olfactory receptor pseudogenes (**Figure 2A**).

Next, we investigate whether SVhound is suggesting regions containing repeats that are known to show many structural variants (Mahmoud et al. 2019). For this we analyzed whether the candidate windows harbored repeat elements (Tarailo-Graovac and Chen 2009) or simple tandem repeat elements (Benson 1999). We found that the LINE and LTR repeat families were the most often observed in the candidate windows, with the L1-LINE repeat (Benson 1999) being the most abundant (**Supplementary tables 4.1** and **4.2**). Because simple tandem repeat elements occur frequently in the human genome, we found all but one window overlapped with at least one simple tandem repeat. Furthermore, we investigated whether these ubiquitous elements were present more abundantly within the candidate windows. **Supplementary Figure 7** shows the distribution of the number of simple tandem repeats in the candidate windows and in a random selection of windows for comparison. We performed a two sample T-test of difference in means and a Kolmogorov-Smirnov test of difference in the distribution to compare the distribution of simple repeats in these two sets of windows. Both tests reject the hypothesis of the distribution being different (Kolmogorov-Smirnov test p-value = 0.2378) or the means being different (T-test p-value = 0.314), and thus there is no significant difference in abundance of simple repeats in candidate windows when compared to the rest of the genome.

We further analyzed the proportion of candidate windows that overlap with segmental duplications (Bailey et al. 2002) and found that 101 candidate windows overlap with at least one segmental duplication (**Supplementary table 5**). In fact, we identified several candidate windows that overlapped with more than one segmental duplication.

Next we wondered if SVhound actually only identifies regions with repeats that likely harbor undetected SV. To assess this we analyzed the proportion of the candidate windows overlapping with the “high-confidence” or benchmark regions defined by the Genome in a Bottle Consortium (GIAB, (Zook et al. 2016, 2020)) representing reliable regions for structural variation detections using short reads (e.g. outside of segmental duplications, low mapping quality regions) and thus potential targets for experimental validation. We found that 170 out of the 188 candidate windows overlapped with at least one of the reported regions (**Supplementary table 6**). Therefore, SVhound indeed reports windows with biological significance rather than enriching for artifacts or only regions known to be variable in the genome (e.g. intergenic).

Next, we applied SVhound to identify differences across multiple ethnicities in the 1000 genomes project. We split the 2,504 genomes into their five ethnic groups according to the 1KGP super-population structure (661 African (AFR), 347 Admixed American (AMR), 503 European (EUR), 504 East Asian (EAS), 489 South Asian (SAS)) and extracted the candidate windows by ethnic group repeating the previous analysis for each ethnicity. **Supplementary Figure 8** shows the candidate windows (top 1 % with highest *p*_*new*_) for each of the five ethnic groups. From the collection of all top 1% candidate windows (total number of distinct windows: 468) we investigated those present in a single ethnic group (ethnicity-specific windows) and thus identified potential regions of high polymorphism specific to a particular ethnic group; and those that occurred in all ethnic groups (ubiquitous windows) and thus represent regions of high polymorphism in the all humankind (**Figure 2B**, **Supplementary Table 7**).

We detected 45 (9.62%) ubiquitous windows, whereas 264 (56.41%) windows were ethnicity-specific, which break down as follows: South Asian, 61; African, 60; European, 57; East Asian, 53; Admixed American, 33. Finally, the remaining 159 (33.97%) candidate windows occurred in two to four ethnic groups.

Next, we investigated the role of the genes in the ubiquitous and the ethnicity-specific windows (**Supplementary table 8**). For the genes in the ubiquitous windows, we found enrichment in biological processes also found in the 1KGP full data set (nitrobenzene metabolic process, cellular detoxification of nitrogen compound, xenobiotic catabolic process, interferon-gamma-mediated signaling pathway, antigen processing) (**Supplementary table 9.1**). When analyzing the ethnicity specific windows, we only found gene enrichment in the South Asian ethnic group for 8 biological processes related to keratinization (tissue development, **Supplementary table 9.6**).

Finally, we analyzed if repeat elements overlap with ubiquitous and ethnic specific candidate windows. Here, the L1 (LINE), ERV1 (LTR) and ERVL-MaLR (LTR) repeats were the most abundant among both ubiquitous and ethnic specific candidate windows (**Supplementary table 10.1**). Next, when analyzing the repeat elements present in a single ethnic group, LTR Gypsy-like is an example that overlaps with the ethnicity specific windows of the African population (Havecker et al. 2004). Similarly, an ERVL-like (LRT) repeat is restricted to ethnicity specific windows for European population, the TcMar-Tc2 (DNA repeat) was found in ethnicity specific windows for the Admixed American population and Satellite-telo in the South Asian population (**Supplementary table 10.2**).

### Identification of polymorphic candidate regions across 19,652 human genomes in the USA

To extend our work further, we applied SVhound to detect regions with undetected SVs in 19,652 genomes of US residents (CCDG data) that include 8,969 European-American, 8,099 Hispanic or Latino-American and 2,584 African-American genomes (Sedlazeck et al. 2020). Again, we considered as candidate windows those representing 1% with the highest probability of detecting a new SV-alleles (*p*_*new*_ ≥ 5. 7%). **Figure 3** shows the distribution of the probabilities to detect a new SV-allele when splitting the genomes in 23,554 windows, highlighting in red the 236 the candidate windows.

**Figure 3.**
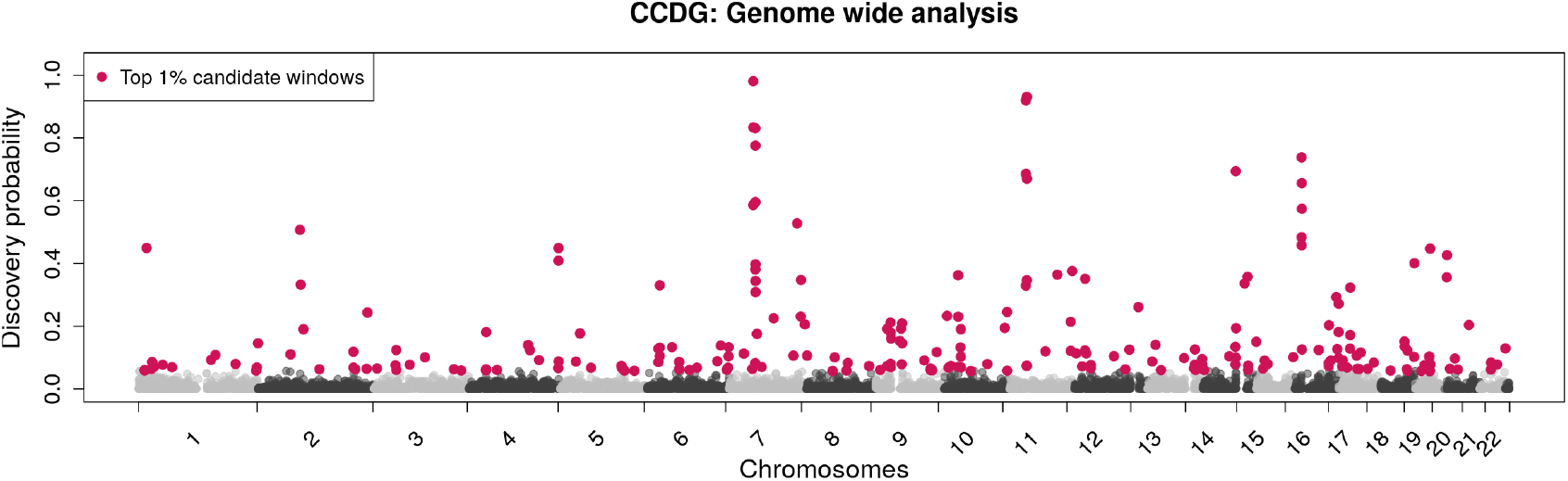
Genome wide analysis of the CCDG data set. Red dots display the top 1% candidate windows (236) along the 22 autosomes of the human genome (hg38), the gray (alternating shades by chromosome) show the discovery probabilities for the remaining windows.

Next, we used a similar annotation strategy to the 1KGP over the 236 candidate windows. We found 156 candidate windows that overlapped with protein coding genes (344, **Supplementary table 11**), 186 overlapping non coding genetic elements (**Supplementary figure 9**) and 33 windows in intergenic regions. Again, we performed an enrichment analysis with PANTHER using the 344 genes and found gene enrichment for 43 biological processes, all of them related to immune response, e.g. phagocytosis, B cell receptor signaling pathway, Fc-gamma receptor signaling pathway involved in phagocytosis, complement activation, positive regulation of B cell activation, innate immune response (**Supplementary table 12**). Next, we analyzed the repeat elements that lay within the candidate windows (**Supplementary table 13**). We observed an overall increase in the number of repeats overlapping with candidate windows. The LINE and LTR families were found in 98.7% and 89% of the candidate windows, which represent an increase of 45% for the LINE and 24% for the LTR when compared to the 1KGP data. In addition, the DNA repeats were found in 60.17% of the candidate windows, while the SINE elements in 49.15% of them, representing an increase of 49% and 45% respectively.

Next, we analyzed the presence of simple tandem repeats within the candidate windows of the CCDG dataset. Here we found significant differences in the average number and the distribution simple tandem repeats across the 236 candidate windows (T-test p-value < 1.24e-13, Kolmogorov–Smirnov test p-value < 2.2e-16, **Supplementary figure 10**). This result again deviates from our analysis of 1KGP data. These results highlight again that the candidate windows that overlapped with centromeric and pericentromeric regions, which tend to be abundant in highly repetitive sequences (Aldrup-Macdonald and Sullivan 2014) and repeats elements and were likely inaccessible/filtered from the 1KGP dataset.

Finally, we noticed consecutive runs of candidate windows along some genomic regions (**Supplementary table 14**). We found such clusters of candidate windows in chromosomes 7 (cluster size 8), 9 (cluster size 5), 11 (cluster size 5) and 16 (cluster size 6). All clusters were located near pericentromeric regions, which have a high density of simple repeats, satellite repeats, and repeat elements in general (LINE, LTR, etc). These results coincide with the instability of the centromeric and telomeric regions in genome assemblies, which are known to be hard to resolve due to their repetitive nature. Thus, these top scoring candidate genomic regions (average *p*_*new*_ in clusters = 43.24%) are confirming already well known to be highly variable.

We then focused on segmental duplications overlapping candidate windows. Here, we observed a slight increase in the number of candidate windows overlapping with a segmental duplication (64.4%) when compared to the 1KGP (53.7%) (**Supplementary table 15**). We identified the candidate windows that overlapped with the GIAB high confidence regions that exclude regions where short reads cannot reliably identify SV. Overall, 86% (203) of candidate windows overlapped with these “high-confidence” regions and thus indicate that reliable SV calling can be achieved in such regions (Zook et al. 2020). (**Supplementary table 16**).

Finally, we compared the results of the two independent human datasets, (1KGP, CCDG) that we analyzed with SVhound to examine the similarities in the prediction. Surprisingly, we found only 26 genes present in candidate windows of both the 1KGP and CCDG data sets, representing approx 5% of the 522 genes associated with at least one of the candidate windows from the 1KGP or CCDG data (**Supplementary table 17**). This small intersection may be related to the fact that the CCDG dataset focuses on the US population while the 1KGP dataset comprises 26 different ethnicities (1000 Genomes Project Consortium et al. 2015), coupled with the difference in number of candidate windows (188 in the 1KGP dataset to 236 in the CCDG dataset, see **Supplementary figure 12**).

### Identification of SV and further polymorphic candidate regions across 150 Rhesus Macaques

To provide a novel test of its utility, we applied SVhound to whole genome sequences from the rhesus macaques (*Macaca mulatta*), a widely used primate model of human disease that has not been well studied with respect to SV (Brasó-Vives et al. 2020; Thomas et al. 2020). For this we created a novel catalog of SV for rhesus macaques by comparing 150 genomes to the newly established reference Mmul_10 (see methods, (Warren et al. 2020)). We identified SVs among the genomes of these 150 rhesus macaques that came from several US research colonies (see methods for details). The largest proportion of SVs were deletions (45.84%) followed by insertions (36.88%), inversions (11.45%) and tandem duplications (5.82%) (**Supplementary table 18.1 and 18.2**). This follows roughly the distribution expected from human SV datasets (Mahmoud et al. 2019). Interestingly, we found a high number of SVs on chromosomes 19 (**Supplementary table 18.3**). Chromosome 19 includes tandem repeats of olfactory receptors, KIR (killer cell immunoglobulin-like receptor) loci and other immunology genes and was previously shown to have a higher rate of both CNV and SNV polymorphism than other macaque chromosomes (Brasó-Vives et al. 2020; Harris et al. 2020). **Figure 4A** shows the minor allele frequency (MAF) spectrum. The MAF spectrum for the genome wide SVs follows the expected exponential distribution, with the majority of the 102,572 SVs (53.7%) exhibiting low frequency (MAF<0.05). We observe 5,946 SV having an MAF >45%, which might be because the reference genome contains an array of low frequency SVs. Interestingly, we noticed a profound peak for Alu insertions (**Figure 4B**) that highlights Alu activity in this species.

**Figure 4.**
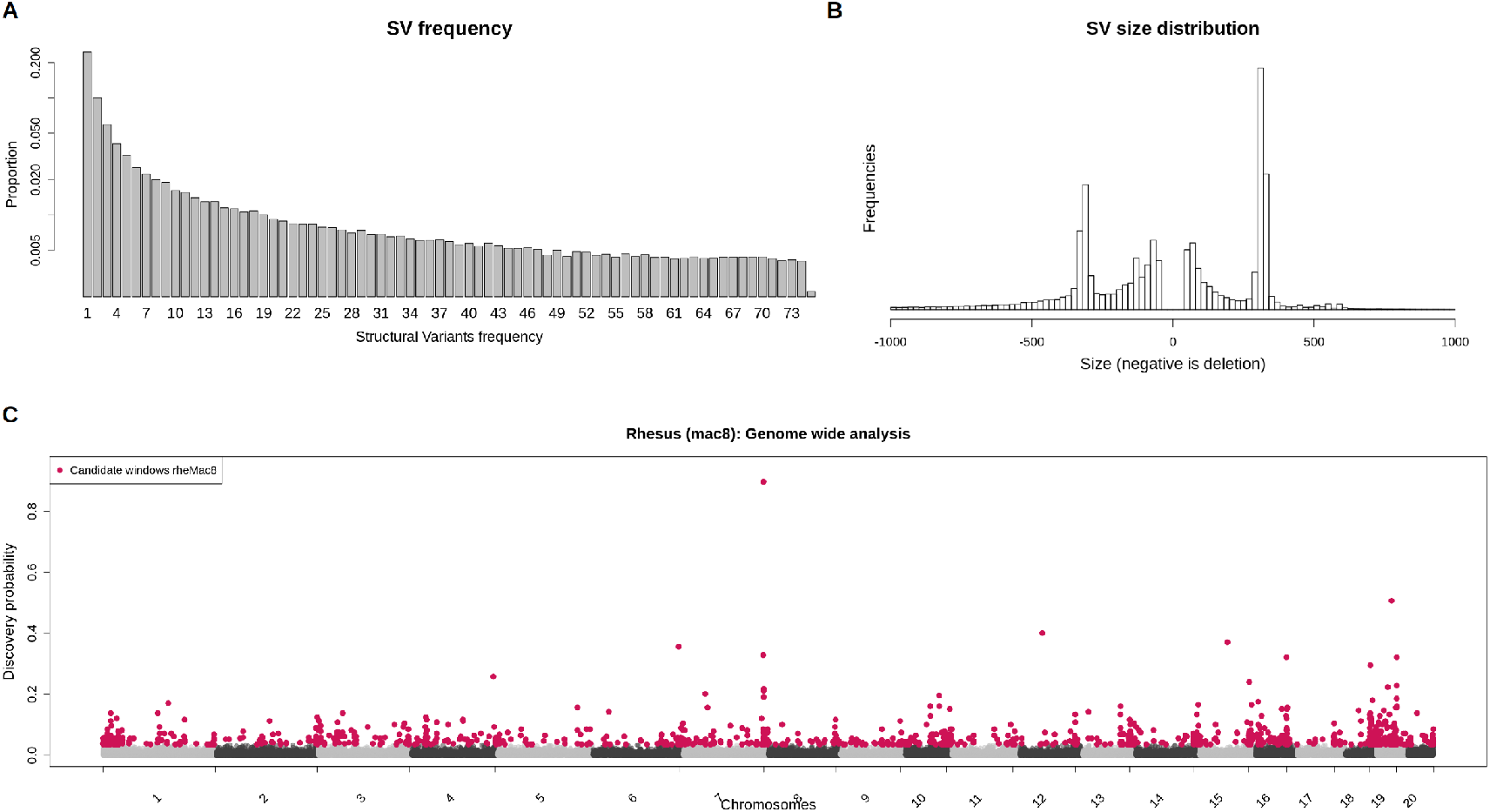
**A)** Frequency distribution of the SV called in 150 rhesus macaque genomes for all SV types. **B)** length distribution of the insertions (positive) and deletions (negative) called in the rhesus macaque genome (truncated at ±1,000 bp, see the full binned table in **Supplementary table 19**). **C)** Genome wide analysis of the rhesus macaque (Macaca mulatta, rheMac8) data set. In red are shown the 1,101 candidate windows (*p*_*new*_ ≥ 3. 57%) along the 20 autosomes of the macaque genome, in gray (alternating shades by chromosome) are shown the rest of the windows.

We applied SVhound to identify candidate regions that may contain undiscovered variation. First, we observed that the rhesus raw data contained a larger number of SVs per window when compared to the human dataset (**Supplementary table 23.1**), even though the number of genomes was an order of magnitude smaller when compared to the 1KGP and two orders of magnitude smaller when compared to the CCDG. **Supplementary Figure 11** shows the candidate windows (top 1% windows with highest *p*_*new*_) for window sizes of 5, 10, 50 and 100 kb. The *p* distribution of the rhesus genome is more spread out relative to the human data. Remarkably the 1% *p*_*new*_ are typically much higher than in humans (Supplementary table 23.2). Starting from 50kb and for larger window sizes, we observed a more widespread distribution of the discovery probabilities *p*_*new*_ which made the selection of candidate windows more difficult. Non-candidate windows had non-negligible *p*_*new*_ values (e.g *p*_*new*_ > 20%). Furthermore, with the larger window sizes the number of regions showing high *p*_*new*_ values increased. In the 100kb window dataset, 24 had the maximum value of *p*_*new*_ dictated by reaching 150 SV-alleles and 1,056 windows had *p*_*new*_ > 50%, whereas in the 10kb window dataset only two windows had a *p*_*new*_ > 50%. With this in mind, we decided to use 10kb windows for the analysis of the rhesus macaque dataset.

We extracted the top 1% candidate windows from the 108,939 10kb windows (*p*_*new*_ ≥ 3. 57%, **Figure 4C**). Then, we extracted 403 annotated rhesus genes that overlap with a candidate window and performed an enrichment analysis with PANTHER (unmapped ID not counted, **Supplementary tables 19**). We found enrichment for divalent metal ion transport related processes and intracellular signal transduction (**Supplementary table 20**).

## Discussion

We developed SVhound to investigate regions along the genome that are likely to harbor undetected SV, exemplifying the method with an analysis of human and rhesus genomes. We were able to demonstrate that these regions harbor genes and are not simply enriched for repeats or intergenic regions along the genome. This indicates their likely importance in evolution and in medicine. SVhound utilizes a sampling scheme approach derived from population genetics (Ewens 1972) to model the SV-allele distribution and to predict genomic regions with high probability of observing new SV-alleles.

SVhound showed a high accuracy over the 1KGP data when assessing its prediction power with a high correlation coefficient across multiple parameters (median correlation across 24 tested parameters = 0.913, best r = 0.993) and slopes close to 1. Apart from the obvious observation that increasing the window size would increase the probability of detecting a new SV-allele (for a 100Mbp window size of course there will be a new SV-allele), we found that small windows (5kb) lead to imprecise predictions, likely due to violations of the model assumptions. Across the human datasets, the method performed well for 100kbp windows (average correlation of 0.894 of 400 evaluations) and even better when considering the windows of 100kb and larger (100kb, 200kb, 500kb, 1mb) where the average correlation was > 0.95 for more than half the evaluations (min. correlation = 0.8189). Remarkably the prediction to find new SV-alleles is sample dependent. The CCDG data with a large sample of 19,000 human genomes exhibited higher *p*_*new*_ values compared to 1KGP (**Supplementary table 21.2**). This difference is resolved if the data processing procedures of the datasets are taken into account. For the CCDG dataset 304,533 SVs were determined, compared to 68,818 SVs for the 1KGP. This difference might reflect the way SVs were called in the 1KGP project, where the majority of genomes had low coverage (3-5x) and likely suffered from a low SV sensitivity, thus leading to an underestimation of the general variability. A conservative SV-calling approach will lead to an underestimation of θ and thus the probability to detect new SVs is also reduced.

The SV-calling procedure in the CCDG project used genomes with a much higher read coverage, thus had more power to detect SVs. These two data sets are hard therefore to compare and clearly shows that SVhound accuracy also relies on the experimental design of the underlying data. The difference might be reduced in the recently posted 1KGP data set where all samples had ~30x coverage (Byrska-Bishop et al. 2021). For rhesus macaques we modified our initial strategy given that, even when we had a smaller cohort (only 150 genomes), a high number of SVs were identified (493,188 SVs), with a different composition (e.g. we identified an abundance of SV especially insertions). Thus, SVhound was run with a smaller window size (10kbp) compared to the human data (100kbp). We have provided guidelines to optimally execute SVhound given different properties of SV cohorts such as the size and the power to detect SV. We anticipate that defining the window size based on the average number of SVs contained in a window may be the path to follow, although more research is needed with a wider number of datasets. Future work will include the automatization of this for SVhound.

Nevertheless, SVhound successfully identified for all three genome projects (1KGP, CCDG, rhesus) genomic regions with a substantial probability to harbor additional SV-alleles. It is noteworthy that SVhound does not require any other annotations than SV coordinates in a region. The candidate regions we found were not confined to well-known regions of high genomic diversity like immune regulatory genes for antigen processing and antigen binding genes (HLA), olfactory genes, regions overlapping repeat elements (LINE, LTR) and regions with an overrepresentation of simple repeat elements (telomeric and pericentromeric regions). Other genomic regions that contained nitrogen related metabolic genes, cellular detoxification related genes and epithelial development genes were also suggested within high probability windows.

It is of course not only interesting which regions SVhound predicts will likely harbor additional not yet observed SV. We can also ask what are the implications? After sequencing hundreds of thousands of genomes, the question might arise whether whole genome sequencing is indeed the most efficient strategy to obtain a more complete set of variations within a particular population of a species. An alternative strategy would be to use a capture design to investigate the identified regions that provide the largest likelihood of containing additional SV-alleles. However, it remains challenging to design these panels for certain regions (e.g. MHC). Nevertheless, it would indeed represent a more efficient strategy to design capture reagents for certain regions and use them to perform targeted sequencing in additional samples to improve the catalog of human population variations. The obvious downside of such an approach is of course that we would likely miss other (rare) SV-alleles in the regions outside of these panels and we don’t know yet if SNV would follow the same trend that we observed for SV. Thus, the challenge remains to obtain a full catalog of common variations across the human population, and also for other important research species. SVhound can assist with prioritizing these regions independently of the organism that is being studied (e.g. non model organism). In addition, SVhound can also indicate that a given population is under-investigated for SV (eg. rhesus data in this manuscript). While this may be obvious given our sample size of 150, it might not be as obvious when the sample size reaches thousands. Here SVhound can again assist in estimating the quality of an SV call set for a given population.

Overall, SVhound shows high prediction accuracy for highlighting regions of the genome where additional SV should be found. This can be resolved either via additional sequencing or improved analysis methods across the data sets in these regions.

## Methods

### Summarization of the structural variants (SV)

We study the genomic variation of a sample of completely sequenced individuals in disjoint fixed windows and analyse each window as follows.

To simplify wording, think of a window as a **locus**, then each distinct SV (particular set of SV present in a given window) is considered as **SV-allele**. For a sample of *n* individuals from this window, we count how often individuals with exactly the same SV in the window occur. With *a*_*i*_ we count the number of different SV-alleles, that occur exactly *i*-times, where 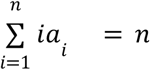 We call *a* = *a*_1_, *a*_2_, …, *a*_n_ SV-occupancy vector. *a*_1_ describes the number of different SV-alleles each occurring exactly once in the sample. If *a*_*n*_ = *n*, then all individuals carry the same SV-allele in the window. Finally 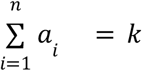 describes the number of different SV-alleles in the window.

We notice that the SV-*occupancy vector* assumes the role of the allele frequency spectrum (AFS) in population genetics (Ewens 1972). However, the AFS is computed for alleles from a gene, whereas the *SV-occupany vector* is computed from the different SV-alleles in a window. Since the potential number of SV-alleles in a window is large, the infinite allele assumption is not severely violated and the celebrated Ewens Sampling Formula (Ewens 1972) that describes the probability to observe a SV-occupancy vector:

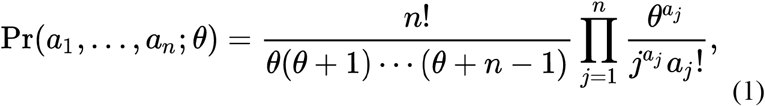

holds, where *θ* is a measure for the genetic diversity of the population. Although Ewens (1972) developed the theory to understand the sampling theory of neutral alleles, we note that the EWS is relevant in very diverse scientific disciplines (see: Harry Crane (2016) The ubiquitous Ewens sampling formula. Statistical Science 31:1-19). Equation (1) and the SV-occupancy vector can be used to compute a maximum likelihood estimator for θ, since this is numerically challenging, we used a simpler approach.

To estimate parameter *θ* based on a sample of *n* individuals, it suffices to apply the method of moment by replacing *E*(*K*), the expected number of SV-alleles by the observed number of alleles *k* and then numerically solve the next equation

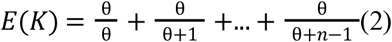

for θ. In fact, 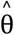 is the maximum likelihood estimate for the data.

Having an estimate, 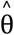, we use this value to compute the “predictive” probability to find a new SV-allele if a new window from an individual is sequenced as:

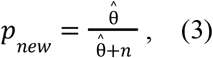

equation 18 in Ewens (Ewens 1972).

Please, note that if θ is small we expect a small number of SV-alleles, a large θ implies that each SV-allele occurs once. However, for such cases to occur θ must be extremely small/large. Finally, notice that *p*_*new*_ = 0 when a single SV-allele is observed (k=1) in the particular window.

To validate SVhound, we partitioned the human genome in non-overlapping windows of size 5, 10, 50, 100, 200, 500, and 1,000 kb. For each window, we randomly re-sampled *n* = 50, 100, 500, 1, 000 individuals without replacement from the 2,504 individuals in the 1,000 human genome project (1000 Genomes Project Consortium et al. 2015) version hg19.This re-sampling was repeated 100 times.

For each subsample, we estimated 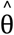 from equation (2) and then estimated the probability to find a new SV-allele, *p*_*new*_, based on equation (3). *p*_*new*_ was subsequently compared to the proportion of individuals that were not in the subsample and that carried SV-alleles not yet detected SV-alleles, that is we computed.

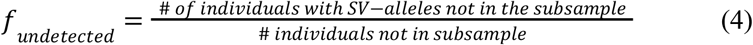

### Identifying SV variability hotspots in the human genomes

We performed a genome-wide analysis to identify genomic regions with a high probability of harboring new SV-alleles. We used two human datasets: a sample of 2,504 individuals for the case of the 1KGP dataset and 19,652 individuals from the Centers for Common Disease Genomics project dataset (Sedlazeck et al. 2020). For both datasets we estimated 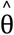 for each window using equation 2 to then calculate the probability of observing a new allele in the next individual using equation 3.

We used a window size of 100kb, which was based on the performance of SVhound prioritizing smaller window sizes. We then selected candidate windows as the 1% windows with the highest probability of detecting a new SV-allele in the next sequenced individual (*p*_*new*_). From these regions we extracted genomic features information from the proper annotation of the human genome (Harrow et al. 2012; Tarailo-Graovac and Chen 2009; Karolchik et al. 2009) (depending on the reference used) to detect what type of genetic elements may be affected.

We performed the enrichment analysis with panther (Mi et al. 2019). We also used data of the position of repeat elements, simple tandem repeats (Benson 1999), segmental duplications (Bailey et al. 2002) and reference “high-confidence” regions form the GIAB project (Zook et al. 2016, 2020).

### Identifying SV variability hotspots in the macaque genomes

We performed a genome-wide analysis to identify genomic regions with a high probability of harboring new SV-alleles. We used a rhesus macaque dataset composed of 150 genomes, for which we estimated 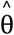 for each window using equation 2 to then calculate the probability of observing a new allele in the next individual using equation 3.

We used a window size of 10 kb, which was based on the performance of SVhound prioritizing smaller window sizes. We then selected candidate windows as the 1% windows with the highest probability of detecting a new SV-allele in the next sequenced individual (*p*_*new*_). From these regions we extracted genomic features information from the rhesus macaque genome annotation Ensembl release 97 (rheMac8, (Yates et al. 2020). We used a list of ortholog genes of rhesus macaque and humans in order to improve the gene enrichment analysis.

### Annotation for the human genome

We used the respective gencode annotation for each of the two versions of the human genomes: genocode 19 for hg19 and genocode 29 for hg38. We complemented the annotation of the genes with the information provided by PANTHER utilizing the Ensemble ID as the gene identifier. We removed all annotated elements (present in gencode) that were not found in Panther (labeled as unmapped IDs).

### Upset plot

All top candidate windows from the five populations (African, American, European, East Esian, South Asian) were pooled. Then for each window its presence/absence was computed for each population (**Supplementary table 7**). Finally for each window the intersection was computed based on the presence/absence binary table. This table was then fed to the upset function of the **UpSetR** library (Conway et al. 2017) according to the reference manual and example.

### Rhesus macaque

We mapped the reads for 150 individuals to the reference of rhesus macaque rheMac8 using bwa mem with default parameter. Subsequently, we identified candidate SVs using Manta (Chen et al. 2016) for each of the bam files separately. Next we computed the region of low mapping quality by extracting reads with MQ<5 and generated a per sample region file by requiring 5 reads of MQ<5 in order to define an interval. The per sample VCF was subsequently filtered by these intervals to account for mapping artifacts and repetitive regions. The resulting VCF files were analyzed and merged using SURVIVOR (Jeffares et al.) merge requiring a SV to be at least 50bp long and up to 1000bp wobble on the start or stop breakpoint.

## DATA ACCESS

Rhesus VCF files (https://github.com/lfpaulin/SVhound) and the R package contain the information of the sources used. 1000genomes VCF file is available at: ftp://ftp.1000genomes.ebi.ac.uk/vol1/ftp/phase3/integrated_sv_map/ALL.wgs.mergedSV.v8.20130502.svs.genotypes.vcf.gz. CCDG VCF file is available over dbVar: nstd160

## ACKNOWLEDGMENTS

This work was supported in part by the US National Institutes of Health (UM1 HG008898 to FJS), DK RNA (UW: W1207-B09) to A.v.H. and NIH grant R24-OD-11173 to J.R.

## References

1000 Genomes Project Consortium, Auton A, Brooks LD, Durbin RM, Garrison EP, Kang HM, Korbel JO, Marchini JL, McCarthy S, McVean GA, et al. 2015. A global reference for human genetic variation. Nature 526: 68–74.

Abel HJ, Larson DE, Chiang C, Das I, Kanchi KL, Layer RM, Neale BM, Salerno WJ, Reeves C, Buyske S, et al. 2018. Mapping and characterization of structural variation in 17,795 deeply sequenced human genomes. Genomics.

Aldrup-Macdonald ME, Sullivan BA. 2014. The past, present, and future of human centromere genomics. Genes 5: 33–50.

Audano PA, Sulovari A, Graves-Lindsay TA, Cantsilieris S, Sorensen M, Welch AE, Dougherty ML, Nelson BJ, Shah A, Dutcher SK, et al. 2019. Characterizing the Major Structural Variant Alleles of the Human Genome. Cell 176: 663–675.e19.

Bailey JA, Gu Z, Clark RA, Reinert K, Samonte RV, Schwartz S, Adams MD, Myers EW, Li PW, Eichler EE. 2002. Recent segmental duplications in the human genome. Science 297: 1003–1007.

Benson G. 1999. Tandem repeats finder: a program to analyze DNA sequences. Nucleic Acids Res 27: 573–580.

Brasó-Vives M, Povolotskaya IS, Hartasánchez DA, Farré X, Fernandez-Callejo M, Raveendran M, Harris RA, Rosene DL, Lorente-Galdos B, Navarro A, et al. 2020. Copy number variants and fixed duplications among 198 rhesus macaques (Macaca mulatta). PLoS Genet 16: e1008742.

Byrska-Bishop M, Evani US, Zhao X, Basile AO, Abel HJ, Regier AA, Corvelo A, Clarke WE, Musunuri R, Nagulapalli K, et al. 2021. High coverage whole genome sequencing *of the expanded 1000 Genomes Project cohort including 602 trios*. Cold Spring Harbor Laboratory 2021.02.06.430068. https://www.biorxiv.org/content/10.1101/2021.02.06.430068v1.abstract (Accessed March 9, 2021).

Chen X, Schulz-Trieglaff O, Shaw R, Barnes B, Schlesinger F, Källberg M, Cox AJ, Kruglyak S, Saunders CT. 2016. Manta: rapid detection of structural variants and indels for germline and cancer sequencing applications. Bioinformatics 32: 1220–1222.

Collins RL, Brand H, Karczewski KJ, Zhao X, Alföldi J, Francioli LC, Khera AV, Lowther C, Gauthier LD, Wang H, et al. 2020. A structural variation reference for medical and population genetics. Nature 581: 444–451.

Conway JR, Lex A, Gehlenborg N. 2017. UpSetR: an R package for the visualization of intersecting sets and their properties. Bioinformatics 33: 2938–2940.

Ebert P, Audano PA, Zhu Q, Rodriguez-Martin B, Porubsky D, Bonder MJ, Sulovari A, Ebler J, Zhou W, Serra Mari R, et al. 2021. Haplotype-resolved diverse human genomes and integrated analysis of structural variation. Science 372. http://dx.doi.org/10.1126/science.abf7117.

Ewens WJ. 1972. The sampling theory of selectively neutral alleles. Theor Popul Biol 3: 87–112.

Goodwin S, McPherson JD, McCombie WR. 2016. Coming of age: ten years of next-generation sequencing technologies. Nat Rev Genet 17: 333–351.

Harris RA, Raveendran M, Worley KC, Rogers J. 2020. Unusual sequence characteristics of human chromosome 19 are conserved across 11 nonhuman primates. BMC Evol Biol 20: 33.

Harrow J, Frankish A, Gonzalez JM, Tapanari E, Diekhans M, Kokocinski F, Aken BL, Barrell D, Zadissa A, Searle S, et al. 2012. GENCODE: the reference human genome annotation for The ENCODE Project. Genome Res 22: 1760–1774.

Havecker ER, Gao X, Voytas DF. 2004. The diversity of LTR retrotransposons. Genome Biol 5: 1–6.

Ho SS, Urban AE, Mills RE. 2020. Structural variation in the sequencing era. Nat Rev Genet 21: 171–189.

Jeffares DC, Jolly C, Hoti M, Speed D, Shaw L, Rallis C, Balloux F, Dessimoz C, Bähler J, Sedlazeck FJ. Transient structural variations have strong effects on quantitative traits and reproductive isolation in fission yeast. http://dx.doi.org/10.1101/047266.

Karolchik D, Hinrichs AS, Kent WJ. 2009. The UCSC Genome Browser. Curr Protoc Bioinformatics Chapter 1: Unit1.4.

Lappalainen T, Scott AJ, Brandt M, Hall IM. 2019. Genomic Analysis in the Age of Human Genome Sequencing. Cell 177: 70–84. http://dx.doi.org/10.1016/j.cell.2019.02.032.

Mahmoud M, Gobet N, Cruz-Dávalos DI, Mounier N, Dessimoz C, Sedlazeck FJ. 2019. Structural variant calling: the long and the short of it. Genome Biol 20: 246.

Mi H, Muruganujan A, Ebert D, Huang X, Thomas PD. 2019. PANTHER version 14: more genomes, a new PANTHER GO-slim and improvements in enrichment analysis tools. Nucleic Acids Res 47: D419–D426.

Sebat J. 2004. Large-Scale Copy Number Polymorphism in the Human Genome. Science 305: 525–528. http://dx.doi.org/10.1126/science.1098918.

Sedlazeck FJ, Yu B, Mansfield AJ, Chen H, Krasheninina O, Tin A, Qi Q, Zarate S, Traynelis JL, Menon V, et al. 2020. Multiethnic catalog of structural variants and their translational impact for disease phenotypes across 19,652 genomes. Genomics 733.

Sudmant PH, Rausch T, Gardner EJ, Handsaker RE, Abyzov A, Huddleston J, Zhang Y, Ye K, Jun G, Fritz MH-Y, et al. 2015. An integrated map of structural variation in 2,504 human genomes. Nature 526: 75–81.

Taliun D, Harris DN, Kessler MD, Carlson J, Szpiech ZA, Torres R, Taliun SAG, Corvelo A, Gogarten SM, Kang HM, et al. 2019. Sequencing of 53,831 diverse genomes from the NHLBI TOPMed Program. Genomics 203.

Tarailo-Graovac M, Chen N. 2009. Using RepeatMasker to identify repetitive elements in genomic sequences. Curr Protoc Bioinformatics Chapter 4: Unit 4.10.

Thomas GWC, Wang RJ, Nguyen J, Harris RA, Raveendran M, Rogers J, Hahn MW. 2020. Origins and long-term patterns of copy-number variation in rhesus macaques. Mol Biol Evol. http://dx.doi.org/10.1093/molbev/msaa303.

Warren WC, Harris RA, Haukness M, Fiddes IT, Murali SC, Fernandes J, Dishuck PC, Storer JM, Raveendran M, Hillier LW, et al. 2020b. Sequence diversity analyses of an improved rhesus macaque genome enhance its biomedical utility. Science 370. http://dx.doi.org/10.1126/science.abc6617.

Wenger AM, Peluso P, Rowell WJ, Chang P-C, Hall RJ, Concepcion GT, Ebler J, Fungtammasan A, Kolesnikov A, Olson ND, et al. 2019. Accurate circular consensus *long-read sequencing improves variant detection and assembly of a human genome*. Nat Biotechnol 37: 1155–1162.

Yates AD, Achuthan P, Akanni W, Allen J, Allen J, Alvarez-Jarreta J, Amode MR, Armean IM, Azov AG, Bennett R, et al. 2020. Ensembl 2020. Nucleic Acids Res 48: D682–D688.

Zook JM, Catoe D, McDaniel J, Vang L, Spies N, Sidow A, Weng Z, Liu Y, Mason CE, Alexander N, et al. 2016. Extensive sequencing of seven human genomes to characterize benchmark reference materials. Sci Data 3: 160025.

Zook JM, Hansen NF, Olson ND, Chapman L, Mullikin JC, Xiao C, Sherry S, Koren S, Phillippy AM, Boutros PC, et al. 2020. A robust benchmark for detection of germline large deletions and insertions. Nat Biotechnol. http://dx.doi.org/10.1038/s41587-020-0538-8.

